# Using unsupervised learning algorithms to identify essential genes associated with SARS-CoV-2 as potential therapeutic targets for COVID-19

**DOI:** 10.1101/2022.05.18.492443

**Authors:** Golnaz Taheri, Mahnaz Habibi

## Abstract

**Motivation:** Severe acute respiratory syndrome coronavirus 2 (SARS-CoV-2) requires the fast discovery of effective treatments to fight this worldwide concern. Several genes associated with the SARS-CoV-2, which are essential for its functionality, pathogenesis, and survival, have been identified. These genes, which play crucial roles in SARS-CoV-2 infection, are considered potential therapeutic targets. Developing drugs against these essential genes to inhibit their regular functions could be a good approach for COVID-19 treatment. Artificial intelligence and machine learning methods provide powerful infrastructures for interpreting and understanding the available data and can assist in finding fast explanations and cures.

**Results:** We propose a method to highlight the essential genes that play crucial roles in SARS-CoV-2 pathogenesis. For this purpose, we define eleven informative topological and biological features for the biological and PPI networks constructed on gene sets that correspond to COVID-19. Then, we use three different unsupervised learning algorithms with different approaches to rank the important genes with respect to our defined informative features. Finally, we present a set of 18 important genes related to COVID-19.

**Availability:** Materials and implementations are available at: https://github.com/MahnazHabibi/Gene_analysis.

**Supplementary information:** Supplementary data are available at *Bioinformatics* online.

## 1 Introduction

As of January 2022, Severe Acute Respiratory Syndrome Corona Virus 2 (SARS-CoV-2), the virus that causes Coronavirus disease 2019 (COVID-19), has infected more than 300 million people worldwide and led to the deaths of more than 5.4 million people. SARS-CoV-2 is a member of the Coronaviridae family of respiratory viruses and it is the third zoonotic coronavirus to emerge in the last two decades. SARS-CoV-2, in comparison to the other two coronaviruses, SARS-CoV (2002) and Middle East respiratory syndrome (MERS)-CoV (2012), has a lower rate of fatality and a higher rate of infection Chen *et al* (2020).

Although there have been thousands of clinical trials, there are no approved medications for COVID-19 yet Thorlund *et al* (2020). However, SARS-CoV-2 has a lower mutation rate than other coronaviruses. On the other hand, high genomic diversity is seen for SARS-CoV-2 both between individual patients and within the same virus class. This diversity enables the virus to adjust to a variety of hosts and circumstances within those hosts and is mostly related to disease development, drug resistance, and treatment results Phan (2020). Therefore, even insignificant but continuous virus alterations and mutations would reduce the efficiency of vaccines or typically used drugs for COVID-19 treatments. Hence, collecting information about the virus’s evolution and pathology will be necessary to control the pandemic situation.

Many researchers are working to identify antiviral drugs and effective vaccines. Therefore, researchers are sharing their findings on SARS-CoV-2’s genome and evolution around the world. Some of these researchers are focusing on finding a therapy with the help of existing drugs using the drug repurposing method as a faster and less expensive approach Aghdam *et al* (2021). Gene analysis is another useful method for drug repurposing and understanding different patients’ responses to the virus. Essential gene analysis can improve the understanding of SARS-CoV-2 data by recognizing the biological pathways of host cells affected by the virus. From the large amount of SARS-CoV-2 related data released, this kind of analysis can help to characterize possible drug targets and drug mechanisms of action Habibi *et al* (2021). As a result, to find an effective treatment, obtaining knowledge from data that characterized the SARS-CoV-2 host infection is a valuable approach.

Several genes associated with the SARS-CoV-2, which are essential for its functionality, pathogenesis, and survival, have been recognized. These genes play crucial roles in SARS-CoV-2 infection and are considered possible therapeutic targets Blanco-Melo *et al* (2020). Ranking important and more relevant genes from all of the COVID-19 associated genes proposed by recent studies will help researchers focus on select sets of genes for further investigation. Developing drugs against these essential genes to inhibit their regular functions and associated physiological pathways could be a good approach for COVID-19 treatment.

In this work, we developed three unsupervised machine learning algorithms to specify important genes, which could help to identify effective COVID-19 treatments. For this purpose, we constructed two biological and Protein-Protein Interaction (PPI) networks corresponding to the COVID-19 related genes. Then, we defined eleven informative topological and biological features for each gene as a node in the network. Finally, we calculated three different scores with respect to our predefined features for each gene with respect to each algorithm. Afterward, we introduced the high score genes in each algorithm with meaningful relationships to COVID-19 as candidate genes for more investigation.

## 2 Materials and methods

In this section, we present a new method to identify essential genes associated with COVID-19 from two inputs: the PPI network and informative biological processes related to COVID-19. In the first step, we calculate four topological features for each protein in the PPI network. We also construct a biological network with respect to informative biological processes related to COVID-19 and calculate five informative features for each protein in the biological network. We also consider two biological features for each protein in the PPI network with respect to COVID-19 pathology. Then, for each protein, we generate a feature matrix *X* = [*x_ij_*]_*m*×*n*_, where *x_ij_* represents the *j*-th feature for the *i*-th protein. In the second step, we use three unsupervised feature selection algorithms (LSFS, RSR, and SPNFSR) to calculate appropriate scores for each feature (*S_j_*). Then, we define the Essentiality Score for each protein (*p_i_*) as follows:

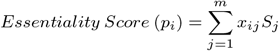

The workflow of the proposed method to identify essential genes is illustrated in Figure 1.

**Fig. 1.**
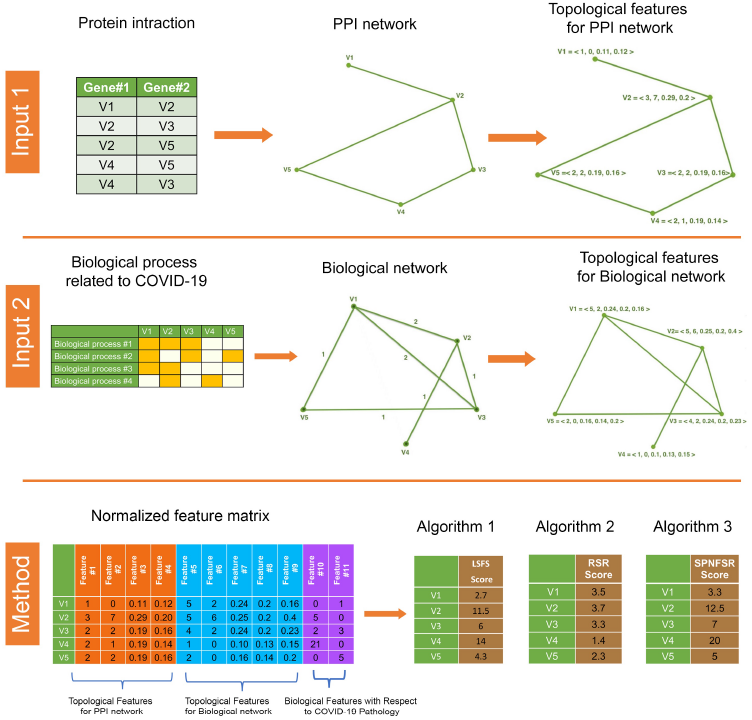
The workflow of the proposed methods.

### 2.1 Datasets

Identifying associated essential genes with disease pathology plays a major role in finding appropriate drugs. Thus, the starting point is to find suitable datasets to extract complete information about proteins and their relationships with COVID-19. For this purpose, we use the PPI network gathered in Habibi *et al* (2021). This dataset contains the physical interactions between proteins that are collected from the Biological General Repository for Interaction Datasets (BioGRID) Chatr-Aryamontri *et al* (2017), Agile Protein Interactomes Data analyzer (APID) Alonso-Lopez *et al* (2019), Homologous interactions (Hint) Patil *et al* (2005), Human Integrated Protein Protein Interaction reference (HIPPIE) Alanis-Lobato. *et al* (2016) and Huri Luck *et al* (2020). All of the proteins in this dataset are mapped to universal protein resource (UniProt) ID UniProt (2019) and those proteins that could not be mapped to a Uniprot ID have been removed. This interactome contains 20,040 proteins and 304,730 interactions. We also use 332 human proteins that interact with the SARS-CoV-2 virus’s proteins that were reported by in Gordon *et al* (2020). These 332 human proteins participate in the 1,374 informative biological processes that are published on the Gene Ontology (GO) Gene Ontology (2019). We denote these informative biological processes as IBP. Among 20,040 proteins, 9,849 proteins participate in the mentioned biological processes.

### 2.2 Informative topological and biological features

In this section, we define informative topological and biological features for each protein in our dataset.

#### 2.2.1 Informative topological features for PPI network

In a topological sense, a PPI network is modeled as an undirected graph *G* =< *V,E* >. Each protein in the PPI network is represented as a node, *v*, and the physical interaction between two proteins (*u* and *v*) is considered as an edge, *uv*. If uv is an edge of graph *G*, a node *u* is the neighbor of node *v*, and the set of neighbors of node *u* is represented by *N*(*u*). A path between *u* and *v* is determined as a sequence of distinct nodes *u* = *u*_0_, *u*_1_,…,*u_n_* = *v* such that *u_i_u*_*i*+1_ is an edge of *G*. The length of a path is equal to the number of edges in this path. The distance between two nodes *u* and *v* is equal to the length of the shortest path between these two nodes, which is denoted by *d*(*u, v*). The following four informative topological features are defined for each node of the PPI network.

1. **Degree**: The number of neighbors of node *u* is defined as degree and denoted by *d*(*u*).
2. **Betweenness**: The betweenness centrality measure of each node *u* on graph *G* is defined as follows:

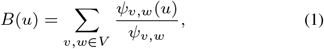

where *ψ_v,w_* denotes the total number of shortest paths between two nodes *v* and *w* and *ψ_v,w_*(*u*) shows the number of shortest paths between two nodes *v* and *w* pass through node *u*.
3. **Pagerank**: Another measure of centrality that is defined for each node *u* is the pagerank. This measure selects the score for each node *u* in the graph as a weighted contribution of all the scores assigned to the node *v* connected to *u* iteratively, as follows:

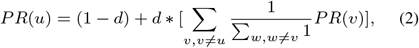

where *d* is a parameter between 0 and 1. *PR*(*u*) is the resulting score vector, whose *i*-th element is the score associated with node *u*. The larger score indicates the importance of the node according to its similarity with the other connected nodes.
4. **Closeness**: The closeness centrality measure for each node, *u*, is defined as follows:

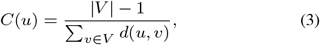

where *d*(*u, v*) is the length of the shortest path between two nodes, *u* and *v*.

#### 2.2.2 Informative topological features for biological network

In this section, we introduce a biological network with respect to 1374 informative biological processes related to COVID-19. This biological network is also modeled as a weighted undirected graph 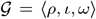. In this graph, each protein that participates in the mentioned biological processes is represented as a node. Two nodes, *u* and *v* are connected through an edge *uv* if two proteins participate in the same biological process. The weight of edge *uv* which is denoted by *ω*(*uv*), is the number of biological processes in which two proteins, *u* and *v*, participate. The length of the path in a weighted graph 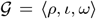, is the sum of the weights of edges encountered when passing through it. The length of a path is equal to the weight of edges in this path. The distance between two nodes *u* and *v* is equal to the weight of the shortest path between these two nodes, which is denoted by *d_ω_*(*u, v*). The following five informative topological features are defined for each node of a weighted biological network.

1. **Weight**: The weight of *u* on weighted graph 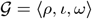, is defined as follows:

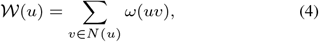
2. **Betweenness**: The betweenness centrality measure of each vertex *u* on graph 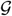 is defined as follows:

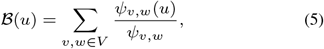

where the shortest path between two nodes, *v* and *w* is determined with respect to the length of the path in the weighted graph.
3. **PageRank**: Another measure of centrality that is defined for each node *u* is the PageRank. This measure selects the score for each node *u* in the graph as a weighted contribution of all the scores assigned to the node *v* connected to *u* iteratively, as follows:

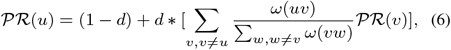

where *d* is a parameter between 0 and 1. 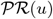 is the resulting score vector, whose *i*-th element is the score associated with node *u*.
4. **Closeness**: The closeness centrality measure for each node, *u*, is defined as follows:

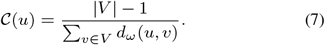
5. **Entropy**: Suppose that *W* = [*w_ij_*] be the weighted matrix correspond to weighted graph 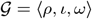 where

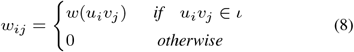

For *j*-th node, we defined *P_j_* as follows:

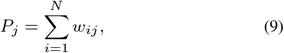

where *N* = |*ρ*|. We also define probability distribution vector *π* =< *P*_1_, *P*_2_, *P_N_* >. Then the entropy of weighted graph is calculated as follows:

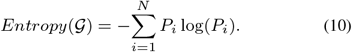

The effect of each node, *u*, on network entropy is defined as follows:

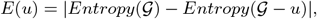

where 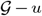 is the weighted network that constructed with respect to removal of node *u* and its connected edges from network.

#### 2.2.3 Informative biological features with respect to COVID-19 pathology

In this section, we also define two biological features for each protein in the PPI network with respect to COVID-19 pathology.

1. For the first feature, we use a set of experimental unapproved drugs in clinical trials for COVID-19 treatment that are available on the Drug Bank Wishart *et al* (2018). This set includes 708 drugs, of which 347 drugs have been studied clinically in more than one clinic. Among these 347 drugs, 213 drugs can target human proteins. This class of drugs is represented by Clinical-Drug. For each protein in the PPI network, the number of drugs approved through this protein is considered the first biological feature related to COVID-19 pathology.
2. For the second feature, we consider the most important signaling pathways related to COVID-19 (NF-*κ*B, Chemokine, Jak-STAT, P53, NOD-like, TNF, CAMP, RAS, Pap1, MAPK, PI3k-Akt, Toll-like(TLR)). For each protein in the PPI network, we calculate the number of these signaling pathways in which the protein participates.

### 2.3 Unsupervised machine learning algorithms

Since the problem of finding the most important set of COVID-19 related genes is still an open question, it can be considered a problem without a response variable or exact answer. Therefore, to find an efficient answer, we used our defined informative features for our constructed COVID-19 related networks. Then, we employed three different unsupervised feature selection algorithms with different approaches to identify an efficient set of genes. It is worth mentioning that, in supervised learning methods, feature selection has been extensively studied. Due to the lack of information about class labels to help the search for relevant knowledge in unsupervised learning methods, selecting features is a significantly more difficult challenge Tomar *et al.* (2020). Suppose *X* = [*x_ij_*]_*m*×*n*_ represents the feature matrix that *x_ij_* represents the *j*-th feature of the *i*-th sample. We assign a feature vector 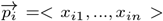 to each sample and define the column matrix *F_j_* = [*x*_1*j*_,…, *x_mj_*]^*T*^ for the *j*-th feature. To find appreciate score for each feature, we use three different unsupervised machine learning algorithms as follows.

#### 2.3.1 Laplacian Score for Feature Selection (LSFS)

Suppose that *S* = [*s_ij_*]_*m*×*m*_ indicates the weighted matrix where 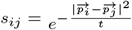 if the euclidean distance between two feature vectors 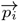 and 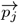 is less than *δ*. Also, suppose that *D* = [*d_i_*] is the diagonal matrix where 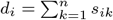 and *L* = *D* – *S* is the Laplacian matrix. The Laplacian Score for each feature, *j*, is calculated as follows:

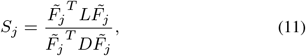

where *J* = [1,1,.., 1]^*T*^ and 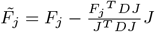.

#### 2.3.2 Non-Convex Regularized Self-Representation (RSR)

Suppose that *W^t^* indicates the weighted matrix and 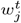 is the *j*-th row of *W^t^*. Let 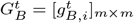 is the diagonal matrix where

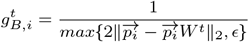

and 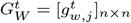 is the diagonal matrix where 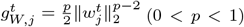. For each 1 ≤ *t* ≤ *N*, the weighted matrix *W*^(*t*+1)^ is calculated iteratively as follows:

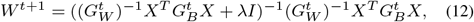

where *I* is the identity matrix and λ > 0. Finally, to compute each feature’s weight using *S_j_* = ||*w^j^*||_2_ (*j* = 1, 2,…, *n*) where *w^j^* denotes the *j*-th row of the weighted matrix *W*.

#### 2.3.3 Structure Preserving Nonnegative Feature Self-Representation (SPNFSR)

Suppose that *S*_*m*×*m*_ indicates the weighted matrix where *S* = (|*S*| + |*S^T^*|)/2 shows the similarity of two feature vectors 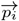 and 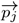. Set two identity matrices *R*_*m*×*m*_, *Q*_*n*×*n*_. Compute matrix *L* = (*I* – *S* – *S^T^* + *SS^T^*) and *M* = *X^T^LX*. Suppose that *M* = *M*+ – *M*^-^ where 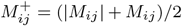 and 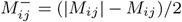. The elements of weighted matrix *W* is calculated iteratively as follows:

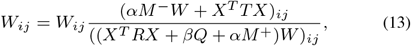

where *α* ≥ 0 and *β* ≥ 0 and two matrices *R* and *Q* as diagonal matrices updated iteratively as follows:

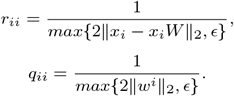

where *ϵ* is a very small constant. Finally, to compute each feature’s weight using *S_j_* = ||*w^j^*||_2_ (*j* = 1, 2,…,*n*) where *w^j^* denotes the *j*-th row of the weighted matrix *W*.

## 3 Results

### 3.1 Evaluation of high score COVID-19 related genes

In this subsection, we studied the 50 top main genes with high scores with respect to three different machine learning algorithms. We evaluated the list of significant diseases and associated pathways related to each of the high score gene sets for each of the algorithms. We also studied the significant modules associated with each of these gene sets.

#### 3.1.1 High score genes selected by LSFS algorithm

The output of the LSFS algorithm is the Laplacian score for each gene. The three genes, TNF, PTGS2, and BCL2, are identified as the three top genes with the highest scores selected by the LSFS algorithm. Studies have shown that TNF could be a key driver of inflammation in patients with severe COVID-19 Zhang *et al* (2020). It could be targeted by existing immunomodulatory therapies. In Li *et al* (2021), the results of molecular docking analysis indicated that niacin showed effective binding capacity in COVID-19 and could help in COVID-19 treatment. One of the important pharmacological targets of niacin in COVID-19 was BCL2 and the other was PTGS2.

Figure 2 also shows that different types of cancer, autoimmune diseases, and diabetes have large numbers of common genes, with the top 50 genes resulting from the LSFS algorithm. Table 1 reports some of the significant disease pathway enrichments identified by the Database for Annotation, Visualization, and Integrated Discovery (DAVID) tools Dennis *et al* (2003). These significant disease pathways like Hepatitis C, Influenza A, Tuberculosis have significant p-values. These pathways contain disease-associated genes that are reported through the LSFS algorithm. From a drug repurposing aspect, effective and most used drugs that target these common genes with selected 50 top genes (for both of the above-mentioned groups of diseases) could be possible COVID-19 treatments. We also studied the important biological processes in these high score gene sets. We used DAVID tool and identified five subsets of biological processes with significant p-values as COVID-19 related modules. Figure 3 illustrates the p-values of each of these modules and the connections between the genes of each of the modules. With the help of the DAVID tool analysis, it was identified that a part of the Fc-epsilon receptor signaling pathway (with a p-value of 1.8 * *E*^-15^) was a submodule in these high score genes. Studies on this module showed that this signaling pathway is followed by the PI3k cascade, which is referred to as the COVID-19 associated pathway Barh *et al* (2021). Studies also have shown that this module is associated with cytokine production in inflammatory diseases Stelzer *et al* (2016). Another identified significant module is a part of the TLR signaling pathway as a MyD88-dependent pathway. In the MyD88-dependent pathway, the MyD88 protein recruits IRAK family proteins. The IRAK4 protein activates TRAF6 and this protein ultimately activates NF-*κ*B resulting in the production of excessive and dangerous inflammatory cytokines in patients with COVID-19 Zhang *et al* (2020).

**Fig. 2.**
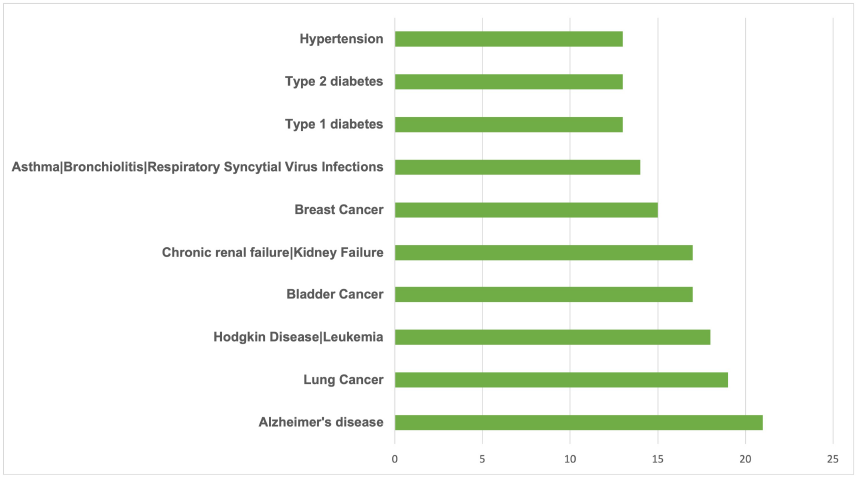
The list of top diseases and number of related disease genes for the LSFS algorithm.

**Table 1.** Top significant disease pathways enrichment identified by DAVID.

| Annotation Cluster 1 (Enrichment Score: : 12.121) |  |  |
| --- | --- | --- |
| Term | Count | P-value |
| hsa05168: Herpes simplex infection | 20 | 4.20E-18 |
| hsa05160: Hepatitis C | 13 | 8.31E-11 |
| Annotation Cluster 2 (Enrichment Score:7.794) |  |  |
| Term | Count | P-value |
| hsa05145: Toxoplasmosis | 19 | 8.34E-21 |
| hsa05160: Hepatitis C | 13 | 8.31E-11 |
| hsa05164: Influenza A | 13 | 1.92E-09 |
| hsa05133: Pertussis | 10 | 1.93E-09 |
| hsa05161: Hepatitis B | 12 | 3.66E-09 |
| hsa05152: Tuberculosis | 12 | 2.99E-08 |

**Fig. 3.**
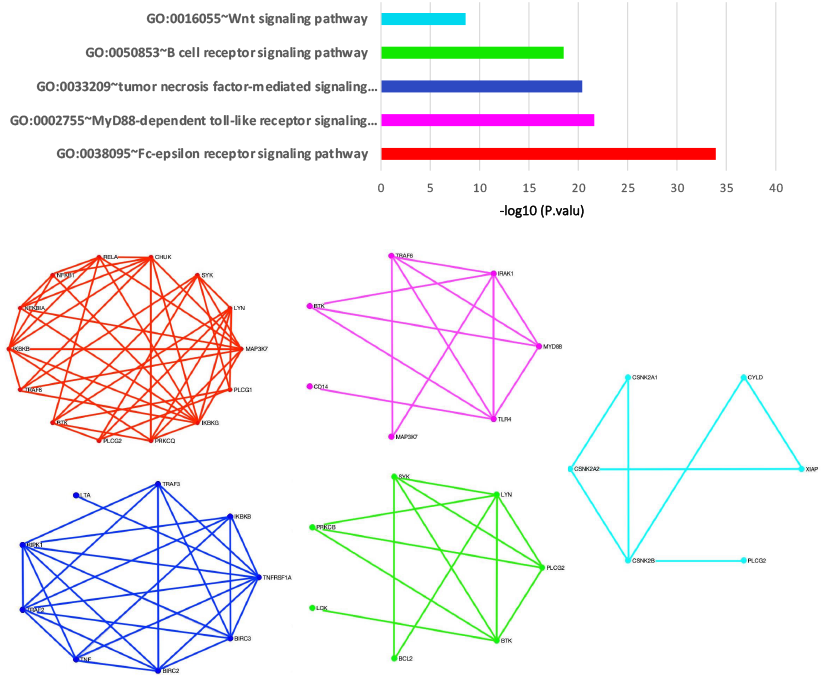
The biological processes with significant p-values for top high score genes through the LSFS algorithm.

#### 3.1.2 High score genes selected by RSR algorithm

The output of the RSR algorithm is the score for each gene. The three genes NTRK1, APP, and ELAVL1, are identified as the three top genes with the highest scores selected by the RSR algorithm. Studies on the NTRK1 gene showed that this gene is associated with the most important symptoms of severe COVID-19, and Fostamatinib, by targeting this gene, has been identified as a therapeutic drug for the control of acute respiratory distress syndrome (ARDS) in COVID-19 patients Wishart *et al* (2018). A recent study showed that the COVID-19 upstream regulators increased APP expression significantly. They revealed that molecular mechanisms of COVID-19 may lead to long-term neurological manifestations resulting from elevated APP expression Camacho *et al* (2021). Another study to prove the value of cellular RNA-binding proteins as therapeutic targets for COVID-19 treatment tested multiple drugs. Their results showed that one of these compounds targeting ELAVL1 caused a meaningful inhibition of SARS-CoV-2 protein production Kamel *et al* (2021).

Figure 4 also shows that different types of cancer and autoimmune diseases have large numbers of common genes, with the top 50 genes resulting from the RSR algorithm. Table 2 reports some of the significant disease pathway enrichments identified by DAVID tool. These significant disease pathways like Hepatitis B and different types of cancers have significant p-values. These pathways contain disease-associated genes that are reported through the RSR algorithm. Therefore, effective, and most used drugs for these diseases that target these common genes with selected 50 top genes could be possible COVID-19 treatments. In Figure 5, we showed the significant submodules associated with these 50 top genes. Figure 5 contains six submodules with a significant p-value from the DAVID tool. We found that these modules have been identified in various studies related to COVID-19 Ghosh *et al* (2021); Hachim *et al* (2020).

**Fig. 4.**
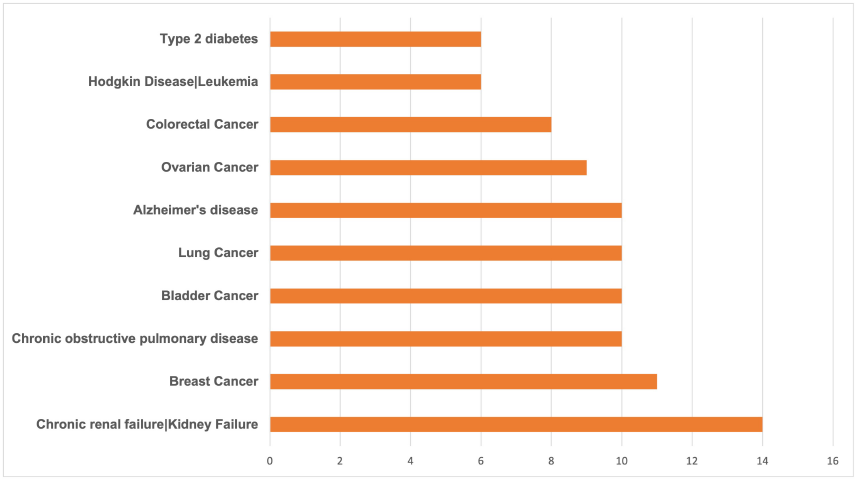
The list of top diseases and number of related disease genes for the RSR algorithm.

**Table 2.** Top significant disease pathways enrichment identified by DAVID.

| Annotation Cluster 1 (Enrichment Score: : 4.211) |  |  |
| --- | --- | --- |
| Term | Count | P-value |
| hsa04110:Cell cycle | 9 | 6.28E-07 |
| hsa05203:Viral carcinogenesis | 8 | 2.09E-04 |
| hsa05161:Hepatitis B | 6 | 0.00176293 |
| Annotation Cluster 2 (Enrichment Score:2.758) |  |  |
| Term | Count | P-value |
| hsa05215:Prostate cancer | 9 | 4.30E-08 |
| hsa05205:Proteoglycans in cancer | 9 | 2.24E-05 |
| hsa05213:Endometrial cancer | 5 | 2.63E-04 |

**Fig. 5.**
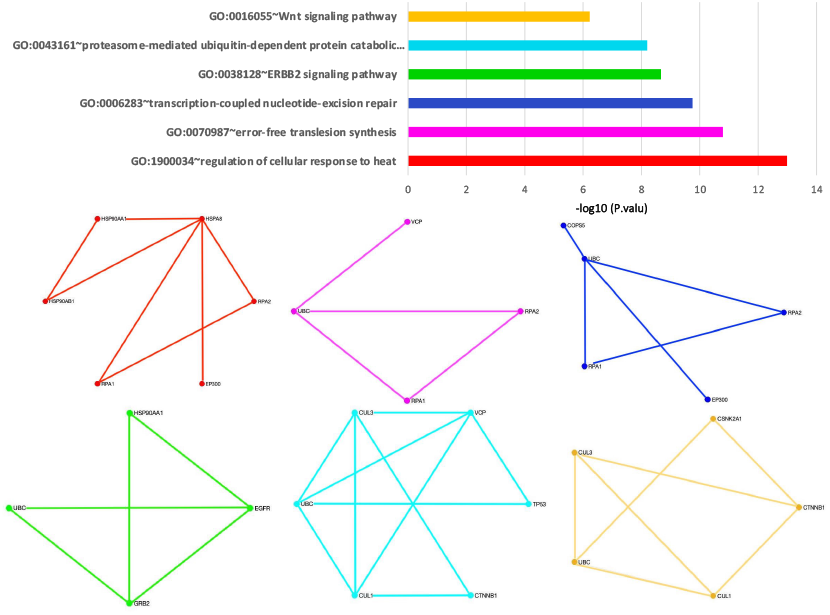
The biological processes with significant p-values for top high score genes through the RSR algorithm.

#### 3.1.3 High score genes selected by SPNFSR algorithm

The output of the SPNFSR algorithm is the score for each gene. The three genes CYP3A4, ABCB1, and CYP2C9, are identified as the three top genes with the highest scores selected by the SPNFSR algorithm. Authors in Kumar *et al* (2021) summarize medication updates for COVID-19 treatment in patients with an inflammatory state and their interactions with drug transporters. They showed CYP3A4, ABCB1, and CYP2C9 could be suitable targets for COVID-19 potential treatments.

Figure 6 also shows that different types of cancer, diabetes and autoimmune diseases have large numbers of common genes, with the top 50 genes resulting from the SPNFSR algorithm. Table 3 reports some of the significant disease pathway enrichments identified by DAVID tool. These significant disease pathways like Influenza A and Rheumatoid arthritis have significant p-values. These pathways contain disease-associated genes that are reported through the SPNFSR algorithm. Therefore, effective, and most used drugs for these diseases that target these common genes with selected 50 top genes could be possible COVID-19 treatments. With the help of the DAVID tool, we identified four significant biological modules. Figure 7 shows the value of each of these modules and the intraction network between them. We found that all of them have been cross-linked with important biological processes or COVID-19 related pathways. Treatment with Ang 1–7 is suggested in several studies. Ang 1–7 decreases the expression of intracellular signaling molecules such as the MAPK family (ERK1/2), which play an essential role in augmenting the inflammatory response Khajah *et al* (2016). Ang 1–7 also inhibits the NF-*κ*B signalings and reduces the expression of Ang II-induced ICAM-1 and VCAM-1. Treatment of COVID-19-affected patients with AT1R blockers (ARBs) may promote the ACE2/Ang 1–7 receptor with the reduction of proinflammatory cytokines and an increment in the level of anti-inflammatory cytokines Gheblawi *et al* (2020).

**Fig. 6.**
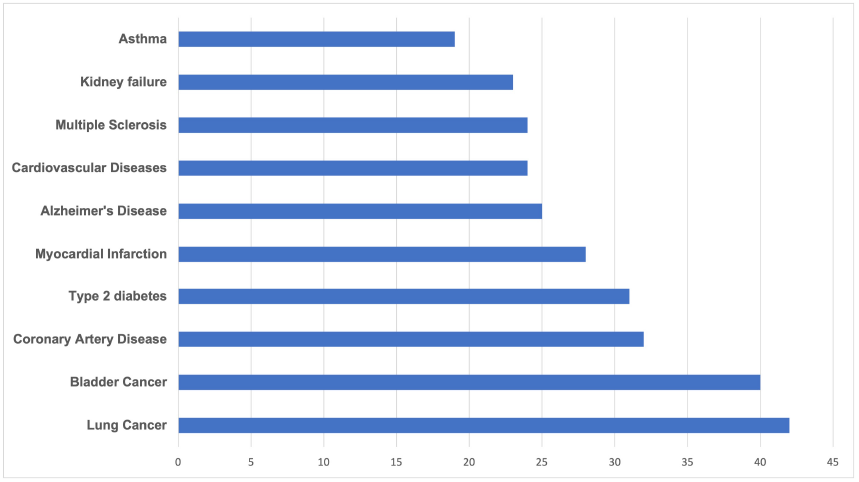
The list of top diseases and number of related disease genes for the SPNFSR algorithm.

**Table 3.** Top significant disease pathways enrichment identified by DAVID.

| Annotation Cluster 1 (Enrichment Score: : 3.057) |  |  |
| --- | --- | --- |
| Term | Count | P-value |
| hsa05212:Pancreatic cancer | 7 | 5,06E-06 |
| hsa04010:MAPK signaling pathway | 11 | 7,22E-06 |
| hsa05160:Hepatitis C | 8 | 3,28E-05 |
| hsa05166:HTLV-I infection | 10 | 5,25E-05 |
| hsa05161:Hepatitis B | 8 | 5,71E-05 |
| Annotation Cluster 2 (Enrichment Score:4.663) |  |  |
| Term | Count | P-value |
| hsa05164:Influenza A | 11 | 2,36E-07 |
| hsa05133:Pertussis | 8 | 7,10E-07 |
| hsa05152:Tuberculosis | 9 | 2,57E-05 |
| hsa05160:Hepatitis C | 8 | 3,28E-05 |
| hsa05161:Hepatitis B | 8 | 5,71E-05 |

**Fig. 7.**
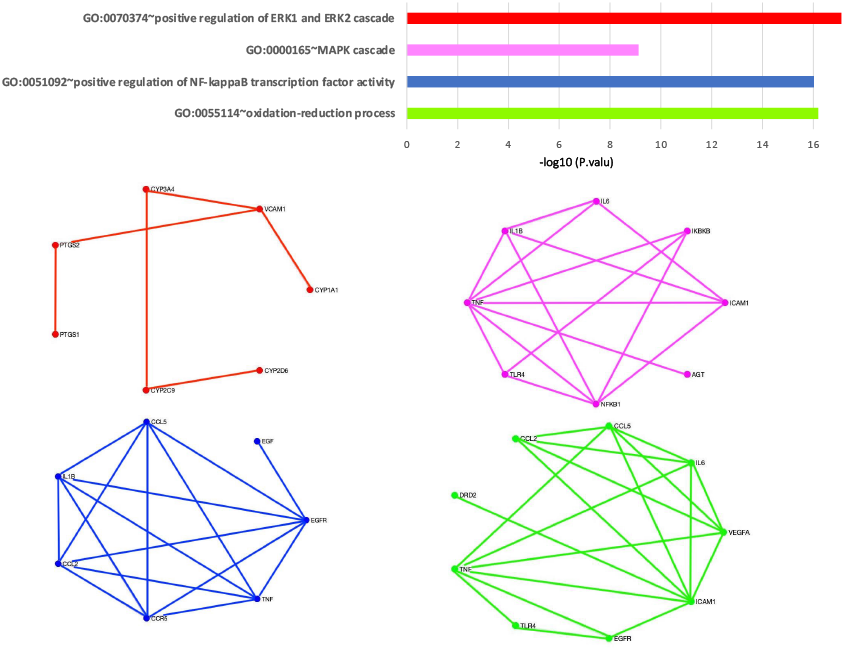
The biological processes with significant p-values for top high score genes through the SPNFSR algorithm.

### 3.2 Evaluate

In this section, we evaluate the results of our proposed algorithms. Our results show that the proposed algorithms provide appropriate models for identifying COVID-19 related genes. We compare and evaluate these three sets of 50 top COVID-19 related genes to suggest a set of the best candidate COVID-19 related genes. We found that 18 genes were identified by at least two of the proposed algorithms. Vascular cell adhesion protein 1 (VCAM1) is reported across all three algorithms. VCAM1 is expressed on inflamed vascular endothelium in inflamed tissue and plays an important role in immune responses UniProt (2019). Also, it makes leukocytes migrate to locations of inflammation UniProt (2019). The systematic analyses showed that the increased expression of VCAM1 is related to COVID-19 disease severity and may contribute to coagulation dysfunction Tong *et al* (2020). Table 4 shows the complete list of these 18 genes and detailed information about their corresponding functions. In Table 4, the genes that have been confirmed in Stelzer *et al* (2016) to be associated with SARS-CoV-2 are shown in bold.

**Table 4.**
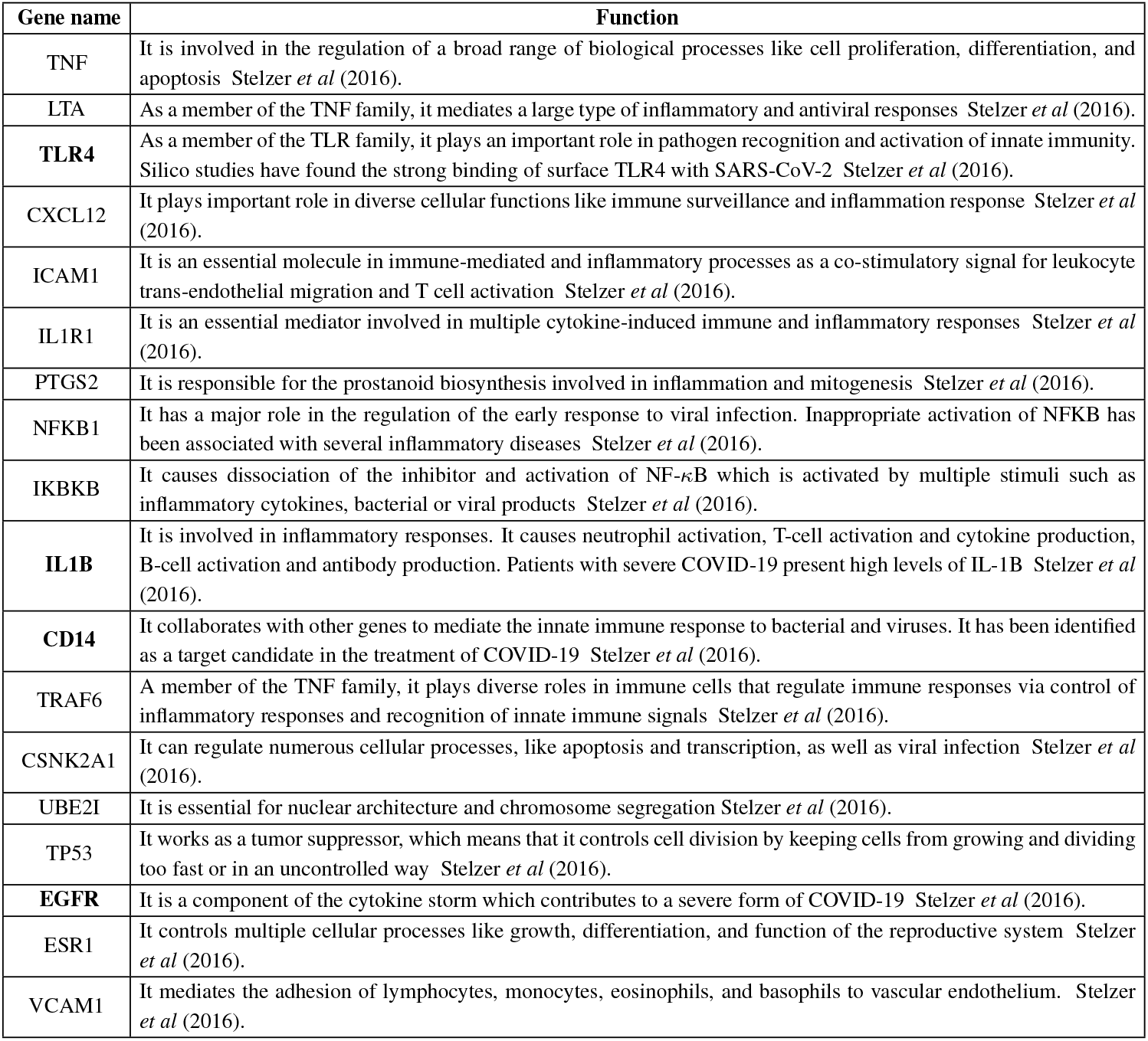
The list of shared genes were identified by at least two of the proposed algorithms and their functions.

## 4 Discussion

The scientific community is trying to find new therapies for COVID-19. For this purpose, researchers face a major challenge in identifying the fewest and most important COVID-19 related genes that could be used as potential drug targets. Numerous studies have been carried out to discover a suitable group of genes associated with COVID-19, and the results of these studies include a long list of genes, each of which could be important. It could be possible to identify effective drug targets by prioritizing these genes based on their topological and biological properties. We presented three machine learning algorithms (LSFS, RSR, SPNFSR) to prioritize COVID-19 related genes and organize these genes. The newly introduced algorithms are based on the feature selection method.

In the first part of this work, we defined 11 biological and topological features for each gene. The first four features, based on the centrality measure of each gene in the PPI network, are introduced as the topological features of the gene. We also built a COVID-19 related biological network. This network was a weighted network that fitted into a set of biological processes containing 332 proteins that were targeted by the virus. In this biological network, we have presented five features according to the topological characteristics of each gene as another measure for each gene. We also defined two other features for each gene in the PPI network. The first one was based on the number of drugs from the Clinical-Drug group that targeted the gene. The second one was based on the number of COVID-19 related signaling pathways that contain the gene. Then, with the help of three unsupervised machine learning algorithms, we assigned a score to these features. We assigned a score to each gene with the help of topological and biological features of each gene and the value of each feature. We prioritized the set of genes based on these scores. In the result part of this work, we looked at the three high-scoring genes in each algorithm and discovered a direct link between these genes and COVID-19. We also evaluated the 50 top high-scoring genes of each algorithm with different measures. In the first measure, we evaluated the common genes between the list of 50 genes and disease genes for each algorithm. Our results show that these genes have the most in common with various types of cancer, diabetes, and autoimmune diseases. As another measure, we reported some of the significant disease pathways like Hepatitis C, Influenza A, and Tuberculosis with significant p-values that contain disease-associated genes that have a lot in common with the list of 50 high-scoring genes. We also studied the biologically significant processes associated with these 50 high-scoring genes. We identified critical modules such as MyD88, Wnt, and MAPK, which have been linked to the SARS-CoV-2 in multiple studies. Finally, we presented a list of 18 genes that have been identified as top genes by at least two of our algorithms. These 18 genes could be targeted by some drugs like Abivertinib, chloroquine, and acetylcysteine that are approved as COVID-19 drugs. In Table 4 we gave some information about these 18 genes. From the information in this table, we found that 4 of these genes have confirmed associations with COVID-19. It can be concluded that these 18 genes could be good candidates as drug targets for further research in clinical trials for possible COVID-19 treatment.

## 5 Conclusion

The spread of COVID-19 and the ways to deal with it have become a major challenge for people around the world. Advanced methods in bioinformatics techniques and computational biology are effective approaches to deal with this problem. In the current situation, the use of these techniques in data analysis and screening with high throughput is a powerful tool. One of the most effective tools in bioinformatics, from the identification of related genes to the design of appropriate drugs for COVID-19, is artificial intelligence techniques and especially machine learning methods. Relying on machine learning algorithms, this paper attempts to find the appropriate set of COVID-19 related genes and prioritize these genes. The presented results show a major success of the machine learning approach against SRAS-CoV-2.

## References

Chen, Y. et al (2020) Emerging coronaviruses: genome structure, replication, and pathogenesis, J of med viro, 92, 418–423.

Thorlund, K. et al (2020) A real-time dashboard of clinical trials for COVID-19, The Lan Digi Health, 2, e286–e287.

Phan, T. (2020) Genetic diversity and evolution of SARS-CoV-2, Infe, gen and evolut, 81, 104260.

Aghdam, R. et al (2021) Using informative features in machine learning based method for COVID-19 drug repurposing, J of cheminfo, 13, 1–14.

Habibi, M. et al (2021) Topological network based drug repurposing for coronavirus 2019, Plos one, 16, e0255270.

Blanco-Melo, D. et al (2020) Imbalanced host response to SARS-CoV-2 drives development of COVID-19, Cell, 181 1036–1045.

Habibi, M. et al (2021) A SARS-CoV-2 (COVID-19) biological network to find targets for drug repurposing., Scie Rep, 11, 1–15.

Chatr-Aryamontri, A. et al (2017) The BioGRID interaction database: 2017 update, Nucl aids rese, 45, D369–D379.

Alonso-Lopez, D. et al (2019) APID database: redefining protein–protein interaction experimental evidences and binary interactomes, Database, 1, 1–8.

Patil, A. et al (2005) HINT: a database of annotated protein-protein interactions and their homologs, Biophysics, 1, 21–24.

Alanis-Lobato. G, et al (2016) HIPPIE v2. 0: enhancing meaningfulness and reliability of protein–protein interaction networks, Nucl aids rese, 1, gkw985.

Luck, K. et al (2020) A reference map of the human binary protein interactome, Nature, 580, 402–408.

UniProt Consortium (2019) UniProt: a worldwide hub of protein knowledge, Nucl aids rese, 47, D506–D515.

Gordon, D. et al (2020) A SARS-CoV-2 protein interaction map reveals targets for drug repurposing, Nature, 583, 459–468.

Gene Ontology Consortium. (2019) The gene ontology resource: 20 years and still GOing strong, Nucl aids rese, 47, D330–D338.

Wishart, D. S. et al (2018) DrugBank 5.0: a major update to the DrugBank database for 2018, Nucl aids rese, 46, D1074–D1082.

Tomar, G. S. et al. (2020) Proceedings of ICSC 2019, Springer Nature., Vol. 1.

Dennis, G. et al (2003) DAVID: database for annotation, visualization, and integrated discovery, Gen bio, 4, 1–11.

Zhang, F. et al (2020) IFN-c and TNF-a drive a CXCL10+CCL2+ macrophage phenotype expanded in severe COVID-19 and other diseases with tissue inflammation, bioRxiv.

Li, R. et al (2021) Network Pharmacology and bioinformatics analyses identify intersection genes of niacin and COVID-19 as potential therapeutic targets, Brief in bioinfo, 22, 1279–1290.

Camacho, R. C. et al (2021) Is Fostamatinib a possible drug for COVID-19?-A computational study.Network Meta-analysis on the Changes of Amyloid Precursor Protein Expression Following SARS-CoV-2 Infection, J of Neur Pharm, 16, 1–14.

Kamel, W et al (2021) Global analysis of protein-RNA interactions in SARS-CoV-2 infected cells reveals key regulators of infection, J of The Instit of Eng, 2, 1–10.

Ghosh, M. et al (2021) Finding Prediction of Interaction Between SARS-CoV-2 and Human Protein: A Data-Driven Approach, Molec Cell, 81, 2851–2867.

Hachim, M. Y. et al (2020) Molecular basis of cardiac and vascular injuries associated with COVID-19, Front in cardio med, 7, 1–15.

Kumar, D. et al (2021) Disease-drug and drug-drug interaction in COVID-19: risk and assessment, Biom & Pharma, 1, 111642.

Barh, D. et al (2021) Predicting COVID-19—Comorbidity Pathway Crosstalk-Based Targets and Drugs: Towards Personalized COVID-19 Management, Biomed, 9, 556.

Tong, M. et al (2020) Elevated expression of serum endothelial cell adhesion molecules in COVID-19 patients, J of infe dise, 222, 894–898.

Khajah, M. A. et al (2016) Anti-inflammatory action of angiotensin 1-7 in experimental colitis, PLoS One, 11, e0150861.

Gheblawi, M. et al (2020) Angiotensin-converting enzyme 2: SARS-CoV-2 receptor and regulator of the renin-angiotensin system: celebrating the 20th anniversary of the discovery of ACE2, Circu res, 126, 1456–1474.

Stelzer, G. et al (2016) The GeneCards suite: from gene data mining to disease genome sequence analyses., Curr proto in bioinfo, 54, 1–30.

